# Carpenter Bee Thorax Vibration and Force Generation Inform Pollen Release Mechanisms During Floral Buzzing

**DOI:** 10.1101/2021.09.28.462062

**Authors:** Mark Jankauski, Cailin Casey, Chelsea Heveran, M. Kathryn Busby, Stephen Buchmann

## Abstract

Approximately 10% of flowering plant species conceal their pollen within tube-like poricidal anthers. Bees extract pollen from poricidal anthers via floral buzzing, a behavior during which they apply cyclic forces by biting the anther and rapidly contracting their flight muscles. The success of pollen extraction during floral buzzing relies on the direction and magnitude of the forces applied by the bees, yet these forces and forcing directions have not been previously quantified. In this work, we developed an experiment to simultaneously measure the directional forces and thorax kinematics produced by carpenter bees (*Xylocopa californica)* during defensive buzzing, a behavior regulated by similar physiological mechanisms as floral buzzing. We found that the buzzing frequencies averaged about 130 Hz and were highly variable within individuals. Force amplitudes were on average 170 mN, but at times reached nearly 500 mN. These forces were 30 – 80 times greater than the weight of the bees tested. The two largest forces occurred within a plane formed by the bees’ flight muscles. Force amplitudes were moderately correlated with thorax displacement, velocity and acceleration amplitudes but only weakly correlated with buzzing frequency. Linear models developed through this work provide a mechanism to estimate forces produced during non-flight behaviors based on thorax kinematic measurements in carpenter bees. Based on the buzzing frequencies, individual bee’s capacity to vary buzz frequency and predominant forcing directions, we hypothesize that carpenter bees leverage vibration amplification to increase the deformation of poricidal anthers, and hence the amount of pollen ejected.

## 1. Introduction

Pollination is essential to the reproduction of flowering plants (angiosperms). A large percentage of such plants conceal their pollen in tube-like anthers (called poricidal anthers) that expose pollen to the environment only through narrow holes or pores [1], [2]. Pollen is extracted from poricidal anthers through bees performing a behavior called “floral buzzing” [3]–[9]. During floral buzzing, a bee grips the anther with its mandibles and rapidly contracts its flight muscles. The subsequent vibration dislodges pollen from the anther’s interior locule walls and eventually causes it to be ejected from the anther’s apical pores [10], [11]. Much of the pollen is collected on the bee’s abdomen and legs, where it may eventually be transferred to the reproductive organs of another flower as the bee continues to forage. The amount of pollen released during floral buzzing is positively correlated to the anther’s vibration amplitude and the duration of the bees’ buzz [10], [11]. The bee can increase the anther’s vibration amplitude (and consequently, pollen release) by: (1) increasing the amount of force applied to the anther, (2) exciting the anther near one or more of its resonant frequencies [12], and/or (3) applying forces to the anther in a direction consistent with large vibration amplification [13], [14]. The effectiveness of pollen extraction therefore relies critically on the vibratile forces imparted to the anther by the buzzing bee, specifically the forcing amplitude, frequency, and direction. Nonetheless, the forces generated by floral buzzing bees, and the directions in which these forces act, have not been adequately characterized.

The forces imparted to a poricidal anther from a buzzing bee originate with the action of internal sets of antagonistic fibrillar muscles known as the indirect flight muscles (IFMs) [15]. The large fibrillar flight muscles of a bee include the dorsal-ventral muscles (DVM) and the dorsal-longitudinal muscles (DLM) [16]. The two DLM muscle sets extend nearly parallel to the long axis of the body, centrally located, and attach to the anterior and posterior surfaces of the inner thorax. The two DVM muscle sets extend approximately along the shorter ventral axis of the body and attach to the dorsal and ventral surfaces of the inner thorax. During flight, small thorax deformations caused by muscle contraction are amplified into large flapping via a linkage system called the wing hinge [17].

The IFMs regulate non-flight behaviors as well, including floral buzzing [18] and defensive buzzing, a behavior intended to deter predators and gain release for the bee [19]. Though both governed by the IFMs, thorax kinematics during flight and non-flight behaviors are distinct, in part because the wings are disengaged during the latter [1]. Disengaging the wings lowers the inertia of the wing-thorax system and may cause thorax vibration frequency and/or amplitude to increase. Vibration frequency and velocity amplitudes, for example, are considerably higher during defensive behaviors than they are during flight in bumblebees [19] and stingless bees [20]. Thorax kinematics observed during defensive buzzing and floral buzzing may differ as well, though in some species they are substantially more like one another than they are to the thorax kinematics observed during flight. In *Bombus terrestris*, for example, buzzing frequencies are approximately 250 Hz and 300 Hz during defensive and floral buzzing respectively, compared to around 130 Hz during flight [19]. Similarly, velocity amplitudes are about 200 mm/s and 250 mm/s during defensive and floral buzzing respectively, and 60 mm/s during flight [19].

Given that defensive buzzing and floral buzzing are regulated by the same physiological mechanisms [21], defensive buzzing is often used as a proxy for floral buzzing [22], [23]. Defensive buzzing is easier to study in a controlled laboratory setting because it can be initiated by lightly pinching or agitating the bee [22]–[24]. By contrast, floral buzzing requires the presence of an anther, and the dynamic response of the anther may distort force measurements. Defensive buzzing is not a perfect proxy for floral buzzing, since recent evidence suggests thorax vibration frequency, velocity and acceleration are modestly higher (cf. 20-30%) during floral buzzing compared to defensive buzzing in bumblebees [19]. It is unclear if thorax kinematics differ between floral buzzing and defensive buzzing in other (non-Xylocopa) bee species due to a lack of comparative studies. Nonetheless, most floral buzzing bees have comparable musculature, and the mechanisms governing non-flight behaviors are similar [25]. Within this work, we therefore assume that defensive buzzing may serve as an approximation of floral buzzing in carpenter bees.

Given the motivation, the first objective of this study was to measure the time-resolved, directional force generation of carpenter bees during defensive buzzing. Knowledge of the directional forces generated by buzzing bees, coupled with structural models of poricidal anthers [26], would enable us to predict how buzz-pollinated anthers deform under realistic dynamic loading. This would in turn facilitate the development of mechanical artificial pollination systems that emulate natural bee pollinators. At the same time, a better understanding of force generation in buzz pollinating bees can help us better understand the ecology of pollinators, such as the physical mechanisms underpinning specialist and generalist behavior between bees and the specific flowers they visit for pollen. For example, floral buzzing bees may leverage vibration amplification by specializing on anthers with natural frequencies within their buzzing frequency range. The second objective of this study is to identify what thorax vibratory properties (frequency, velocity amplitude, etc.) are most strongly correlated with force generation. If kinematic properties such as velocity or acceleration amplitude are highly correlated to force amplitude during defensive buzzing, it may be possible to approximate the forces produced during floral buzzing based upon reported kinematic quantities which are more easily measured. This provides an indirect method to estimate floral buzzing forces, which are challenging to ascertain experimentally due to the presence of the anther.

## 2. Materials and Methods

### 2a. Specimen Collection

We used adult female carpenter bees *Xylocopa (Xylocopoides) californica arizonensis* for all experimental studies. Live bees were collected from sites around Pima and Santa Cruz Counties (AZ) on September 27 and 29, 2020; and October 11 and 13, 2020. All subjects were obtained from within their tunnel nests inside one year old Sotol (*Dasylirion wheeleri*) dried inflorescence stalks (Figure 1). In these cases, whole stalks containing nests were removed, entrances sealed with a rope knot, and brought to a lab setting where the stalks were split longitudinally with a beekeeper’s hive tool. Due to this collection method, all adults were most likely newly emerged young adults. Resident bees within were fed a 50% sucrose solution and then transferred to 15 mL plastic Eppendorf tubes with added ventilation holes. A moist piece of cardboard was added to line each Eppendorf tube to physically support the bees and for added moisture. The tubes were then refrigerated at 4°C for 1 – 6 days. Prior to shipment, bees were removed from the refrigerator, warmed to room temperature, and fed a 50% sucrose solution before being repacked into individual tubes. Tubes were insulated with a Blue Ice pack and/or Styrofoam cylinder, all of which was contained within a cardboard cylinder for shipping. Bees were shipped overnight to the site of the subsequent experiments (Montana State University, Bozeman, MT), with all shipments arriving within one week or less of initial field collection. Upon arrival, bees were stored in a refrigerator at 4°C until testing. Prior to testing, bees were removed from the refrigerator and warmed to ambient room temperature for approximately 5-10 minutes. Following testing, bees were sacrificed by placing them in the freezer. All bees were tested within 24 hours of arrival at Montana State University.

**Figure 1.**
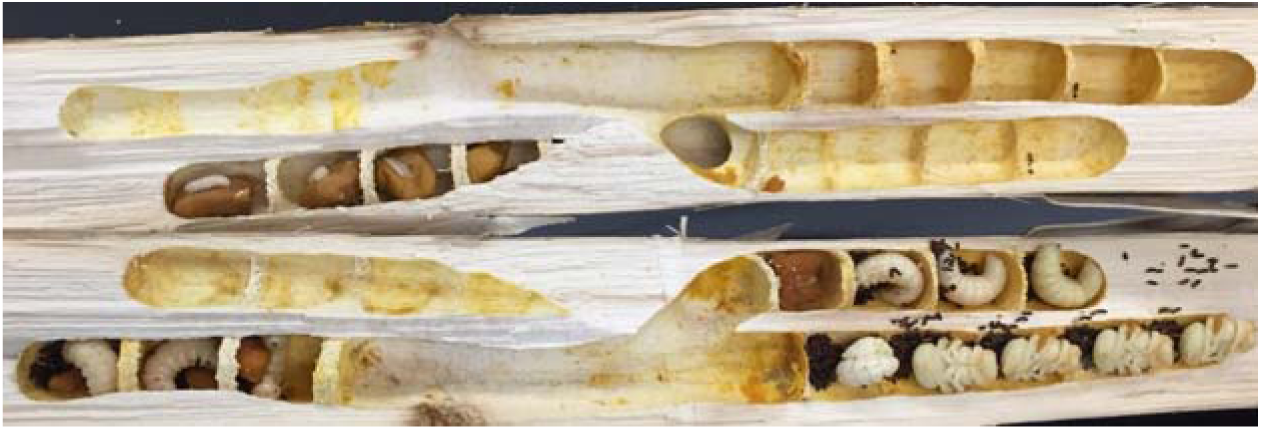
Dried Sotol stalks where carpenter bees were collected. Note the presence of larvae and pupae.

### 2b. Experimental Design

We developed an experiment to measure the directional forces and thoracic vibrations during defensive buzzing in carpenter bees. Forces were measured using a high-sensitivity six axis Nano17-E force-torque transducer (ATI Industrial Automation, Apex, NC, USA) with 3.125 mN force resolution in each axis. We mounted a cylindrical post to the force transducer (Figure 2) that bees were fixed to during experimental trials. The post was 3D printed using a FormLabs Form 2 SLA printer (Formlabs, Somerville, MA, USA) with proprietary tough resin, which has properties similar to Acrylonitrile Butadiene Styrene (ABS). The plastic post had a natural frequency of 715 Hz. Initial testing indicated that bees produced higher-order force harmonics that extended to frequencies past the post’s natural frequency. To identify if post stiffness influenced force generation or thorax vibration, we 3D printed a second post from continuous carbon fiber with 37% triangular infill using a Markforged X7 printer (Markforged, Watertown, MA, USA). The carbon fiber post increased the first resonant frequency of the system to 1400 Hz. During experimental trials, thorax deformation velocity was measured using a non-contact laser vibrometer (Polytec, PSV-400, Hudson, MA, USA) at the thoracic scutum. The scutum had adequate reflectance and did not require treatment with reflectance enhancing agents. We numerically integrated and differentiated to calculate thorax displacement and acceleration, respectively.

**Figure 2.**
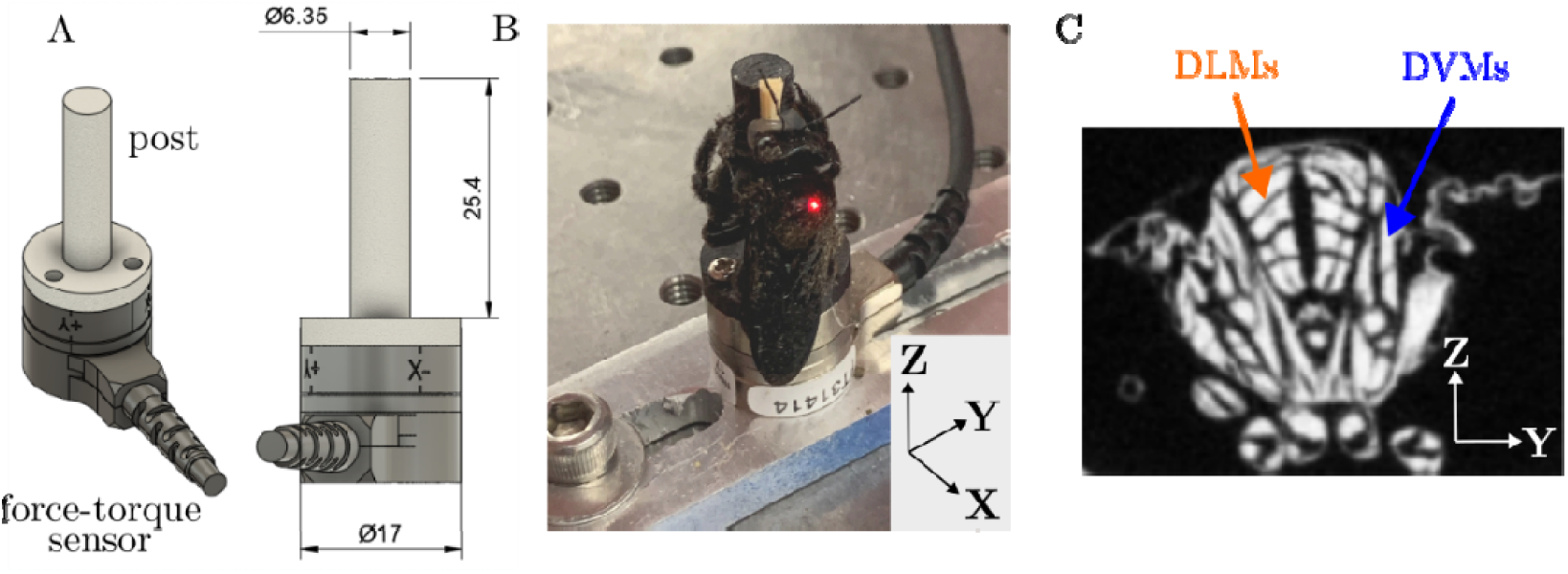
Experimental set-up. (A) CAD model. Dimensions are in millimeters. (B) Carpenter bee mounted to a carbon fiber post. Note the vibrometer laser spot on the dorsal surface of the bee thorax. The cartesian basis shown in the lower left corner defines force directions. The x and z axes most closely align with the insect’s DVM and DLM muscle groups, respectively. (C) MicroCT scan shows a transverse cross section of the insect thorax with flight musculature. The DVM and DLM muscle groups are indicated by arrows. Note that the microCT image shows a honeybee thorax, which has flight musculature that is anatomically similar compared to the carpenter bees considered in this study.

Cold-anesthetized insects were mounted to a wooden toothpick by their mandibles via low temperature hot glue. Glue was extended to the labrum and clypeus to provide additional adhesion so that the bee could not free itself during experimentation. The toothpick was subsequently glued to the post and trimmed to match the post’s length. After mounting, the bees were left to warm up at ambient room temperature. While it is possible that this tethering mechanism influenced the bee’s behavior compared to unconstrained defensive buzzing, the rigid tether at the mandibles was necessary to ensure force measurements were not distorted before reaching the force transducer.

We then lightly pinched (with metal forceps) the bee’s hind tarsus or tibia to initiate defensive buzzing and repeated when the bee quiesced. We recorded at least 2 minutes of buzzing per insect. Five subjects each were tested on the plastic post (subjects 1-5 hereafter) and five subjects were tested on the carbon fiber post (subjects 6-10 hereafter). In total, ten individuals were tested. The bee’s DVM and DLM flight muscles were approximately aligned with the force transducer’s x and z axes respectively, and the transducer’s y-axis runs medially-laterally (Figure 2). A representative video of a defensively buzzing bee is included in the manuscript supplementary material.

The bees were free to move their body during experimentation, which posed challenges for data collection. The orientation of the head is fixed, but the bee can twist its thorax and abdomen or reposition its legs as it attempts to free itself. While muscle forces are transmitted to the post primarily through a rigid connection at the bee’s mandibles, a small force component may be transmitted through the bee’s legs as well. The motion of the bees’ body may influence force directionality. We consequently discarded trials where the vibrometer signal exceeded a prescribed noise level. High vibrometer noise is associated with poor optical reflectance, which occurs when the bee is moving and the scutum plane is not normal with respect to the measurement laser. Numeric criteria for data exclusion are described in the following section.

### 2c. Data Analysis

Time-series force and velocity data were acquired at a sampling rate of 10 kHz and were post-processed in MATLAB 2020B. We applied a bandpass filter with a low frequency cut-off of 10 Hz and a high frequency cut-off of 2000 Hz. Each trial was then divided into equal intervals of 1024 samples, where each interval overlaps 50% with the previous interval. We established two criteria that each interval must satisfy for inclusion in tabulated data. First, the root-mean-squared (RMS) value of the force recorded in the x-axis must exceed 10 mN. This criterion is designed to exclude periods of quiescence. Second, the RMS value of the unfiltered velocity magnitude between 2.5 – 5 kHz must be less than 0.5 mm/s p-p. This criterion excludes intervals in which the optical reflectance is low, which indicates a high angle of incidence between the scutum and vibrometer laser. On average, 514 intervals per bee met the inclusion criterion (min = 35, max = 1336, std = 420). Individuals 1-10 had 711, 244, 390, 1336, 136, 379, 35, 100, 693 and 1122 measurement intervals, respectively.

For each valid interval, we tabulated maximum values for forces in the x, y and z directions as well as thorax integument velocity. To determine thorax oscillation frequency, we applied a Hamming window to each velocity time series and subsequently converted the time series to the frequency domain via a Fast Fourier Transform (FFT). The Hamming window is designed to reduce spectral leakage. We used MATLAB to identify velocity magnitude peaks that exceeded 1 mm/s p-p and that were spaced at least 20 Hz from adjacent peaks. In most cases, the velocity magnitude occurred at some fundamental frequency and several integer harmonics thereof; we reported only the dominant frequency where most of the vibrational energy occurs in the present research. A representative data set is shown in Figure 3.

**Figure 3.**
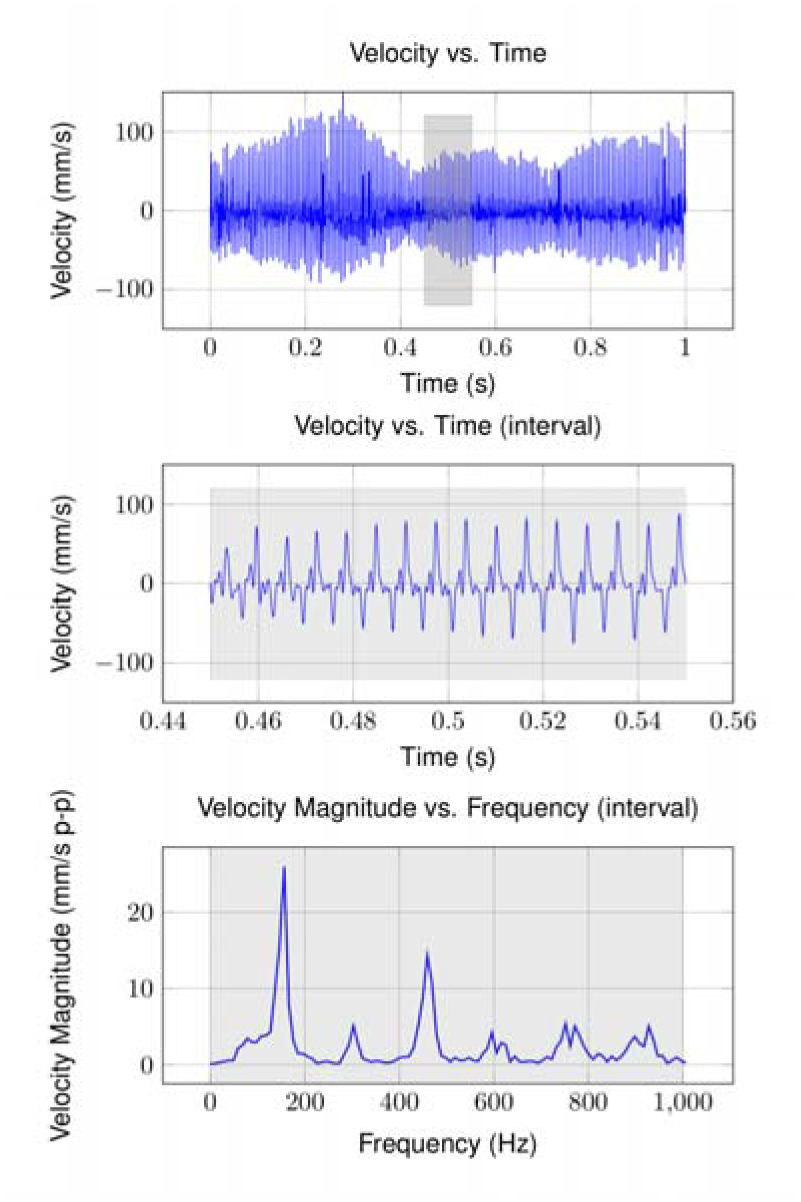
Representative velocity measurements. (Top) Time series data over one second of continuous buzzing. (Middle) Individual interval from which thorax kinematics and force values are calculate. (Bottom) FFT magnitude of interval data. Thorax oscillation frequency is estimated from each velocity interval FFT. Note the frequency harmonics superimposed on the fundamental oscillation frequency.

### 2d. Statistics

To test whether measurements were affected by the post type, mixed model ANOVA was performed using bee (i.e., individual specimen ID) as a random effect and post as a fixed effect. Mixed model ANOVA also tested whether force depends on direction (*F*_*x*_, *F*_*y*_, *F*_*z*_), post, or the interaction between these factors. This test was conducted on data averaged over each interval. For all models, significance was set *a priori* to p < 0.05. Responses were natural-log transformed if necessary to satisfy ANOVA assumptions of residual normality and homoscedasticity. All statistical analyses were performed using Minitab (v.18).

Then, we performed linear regression to fit models relating peak force to thorax vibration frequency, displacement, velocity, and acceleration. We performed this regression on measurement intervals for the entire tested population as well as for individual bees. When regressing peak force against thorax kinematic amplitudes, we divided each measurement interval into four subintervals. Subdividing intervals improved time resolution of force and kinematic measurements and the correlations between the two. For each linear model, we determined intercept, slope, and Pearson’s correlation coefficient.

## 3. Results

### 3a. Effect of Post Material

Post material did not significantly influence buzzing frequency, peak forces in any direction, or displacement, velocity, and acceleration amplitudes (p > 0.05 for effect of post for each test) (see supplementary material). As a result, we could average measurements between the carbon and plastic post groups. For both post materials, low magnitude force harmonics extended to frequencies comparable to the system’s resonant frequency. The force magnitude evaluated at frequencies near the system’s resonant frequency will consequently be amplified. The extensive higher-order harmonics indicate that force is applied to the post by the bees in short periodic pulses rather than harmonically. Nonetheless, bees did not excite the post resonance. Force amplitudes therefore reliably indicate the peak forces transmitted to the post by the bees.

### 3b. Force Generation and Directionality

Interval data for buzzing frequency and displacement, velocity, and acceleration amplitudes are shown in Figure 4. Kinematic data averaged across the population are shown in Table 1. There has been limited data reported for the *X. californica* during defensive buzzing, so we compare our findings to kinematics reported for other Hymenoptera in the text that follows.

**Figure 4.**
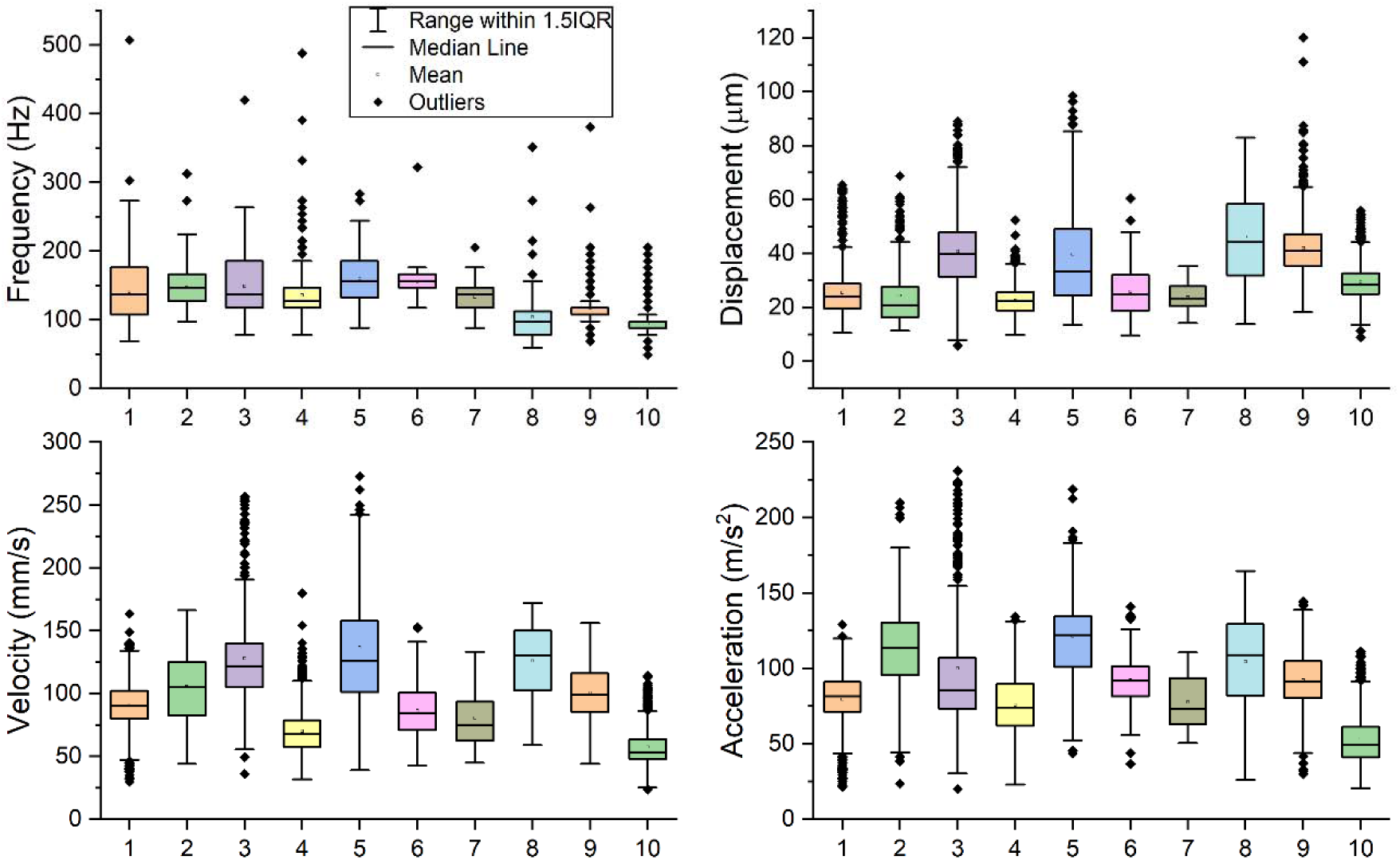
Thorax kinematics (frequency, peak displacement, velocity, acceleration) over all intervals for all carpenter bee individuals (1-10 on the x-axis). Individuals 1-5 were tested on the plastic post, and individuals 6-10 were tested on the carbon fiber post. Individuals 1-10 had 711, 244, 390, 1336, 136, 379, 35, 100, 693 and 1122 measurement intervals, respectively.

**Table 1.** Thorax kinematics (frequency and displacement/velocity/acceleration amplitudes) averaged across the population (n = 10).

| Quantity | Mean | Standard Deviation | Q1 | Q3 |
| --- | --- | --- | --- | --- |
| Frequency (Hz) | 133.5 | 20.8 | 121.3 | 148.0 |
| Displacement ($\mu\text{m}$) | 32.0 | 8.6 | 24.7 | 40.6 |
| Velocity (mm/s) | 98.1 | 24.8 | 82.0 | 120.7 |
| Acceleration ($\text{m/s}^2$) | 91.0 | 19.1 | 78.2 | 103.4 |
| $F_x$ (mN) | 172.3 | 46.9 | 139.9 | 203.3 |
| $F_y$ (mN) | 67.3 | 19.3 | 64.0 | 80.0 |
| $F_z$ (mN) | 115.1 | 36.3 | 83.2 | 133.1 |

The population-averaged 133.5 Hz buzz frequency fell within the 110 – 240 Hz range reported for a wider range of Hymenoptera [21], but was lower than 250 Hz frequency reported for smaller *Bombus terrestris* [19]. Mean displacement (32.0 µm) and velocity amplitudes (98.1 mm/s) were about 27% and 50% lower respectively compared to those measured in *Melipona seminigra* [20], and approximately 85% and 50% lower respectively than those measured in *Bombus terrestris* [19]. Mean acceleration amplitudes (91.0 m/s^2^) occurred within the 60 – 200 m/s^2^ range reported for various Hymenoptera [21]. Kinematics quantities were highly variable across individuals. Similarly large variations in population-averaged frequency and accelerations have been observed in a wide range of hoverflies and bees [21] as well as several species in the bumblebee taxa [27]. Though mean values of thorax vibration frequency and displacement/velocity/acceleration amplitudes were similar between individuals, there were significant variation among these quantities *within* individuals. This suggests that defensive buzzing is a non-stationary behavior and may not be well described by mean values alone.

The forces amplitudes for individual bees are shown in Figure 5. Force data averaged across the population are shown in

**Figure 5.**
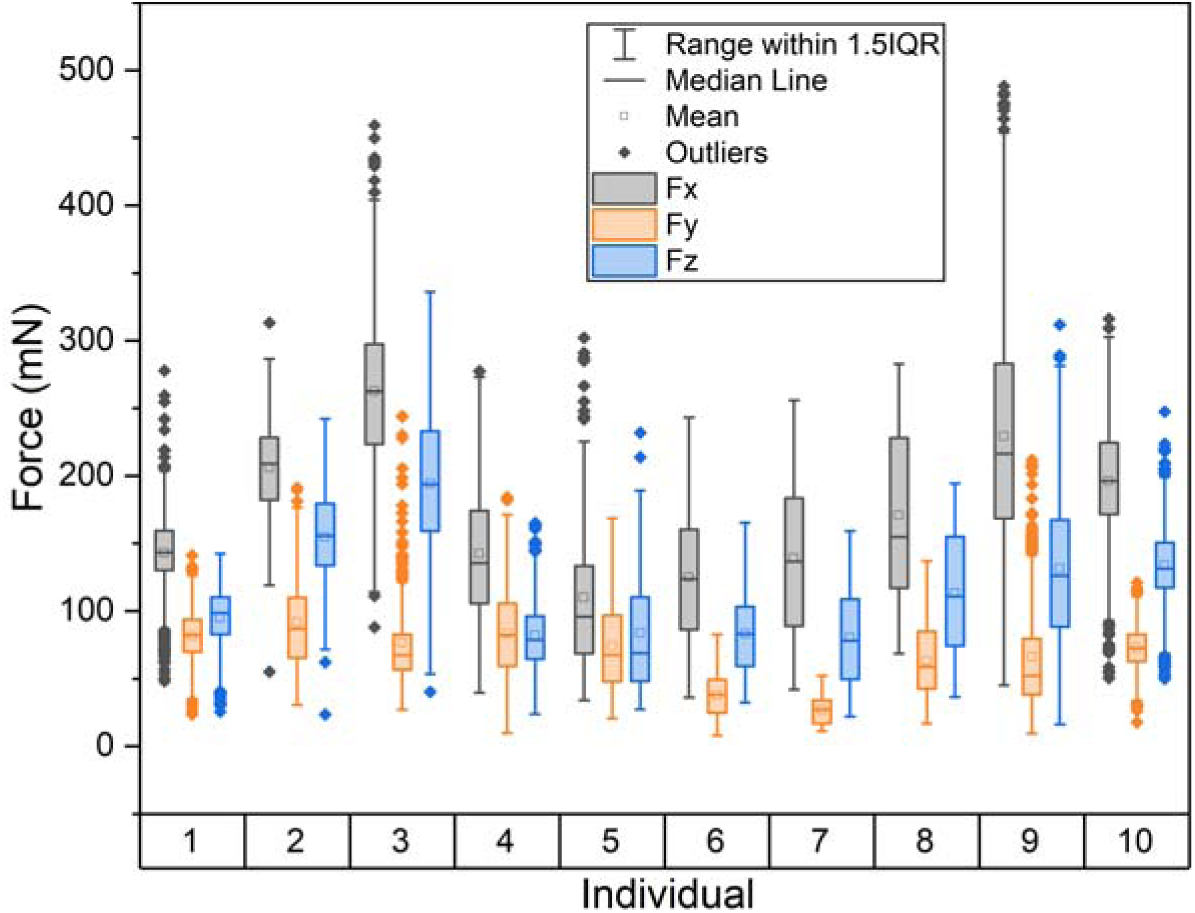
Force amplitudes by direction over all intervals for all individuals. Force amplitudes in the x, y, and z directions are indicated by F_x_, F_y_ and F_z_, respectively. Individuals 1-5 were tested on the plastic post, and individuals 6-10 were tested on the carbon fiber post. Individuals 1-10 had 711, 244, 390, 1336, 136, 379, 35, 100, 693 and 1122 measurement intervals, respectively.

Table 1. Mixed model ANOVA found that forces were not equivalent in x, y, and z directions (p < 0.001, see supplementary material). Post type was not significant and there was not an interaction between post type and force direction (p > 0.05 for both). Post-hoc testing revealed the force in x (172.3 mN) exceeded that in z (115.1 mN), which exceeded that of y (67.4 mN). The x and z directions form a plane containing the bee’s IFMs (Figure 2). Compared to the forces generated in the x direction, forces in the z and y directions were about 32% and 57% lower respectively. Individually, only one bee (subject 4, Figure 5) produced forces in the y direction on the same order as those produced in the z direction. Average forces were highly variable across the population as well as within individuals. To our knowledge, there is no published force data to compare our measurements against. However, reported magnitudes are on the same order as those produced by bumble bee muscle during studies where the muscle is artificially actuated *ex-vivo* [28]. Like thorax vibration frequencies and displacement/velocity/acceleration amplitudes, force amplitudes in all directions were highly variable within individual bees.

### 3c. Correlations Between Thorax Kinematics and Force

Thorax oscillation frequency and displacement/velocity/acceleration amplitudes are shown against peak force in the x-direction (*F*_*x*_) for all measurement intervals for all bees in Figure 6. Across the population, *F*_*x*_ was weakly negatively correlated (r=-0.11) with buzzing frequency and moderately positively correlated with displacement (r=0.41), velocity (r=0.53) and acceleration amplitudes (r=0.51).

**Figure 6.**
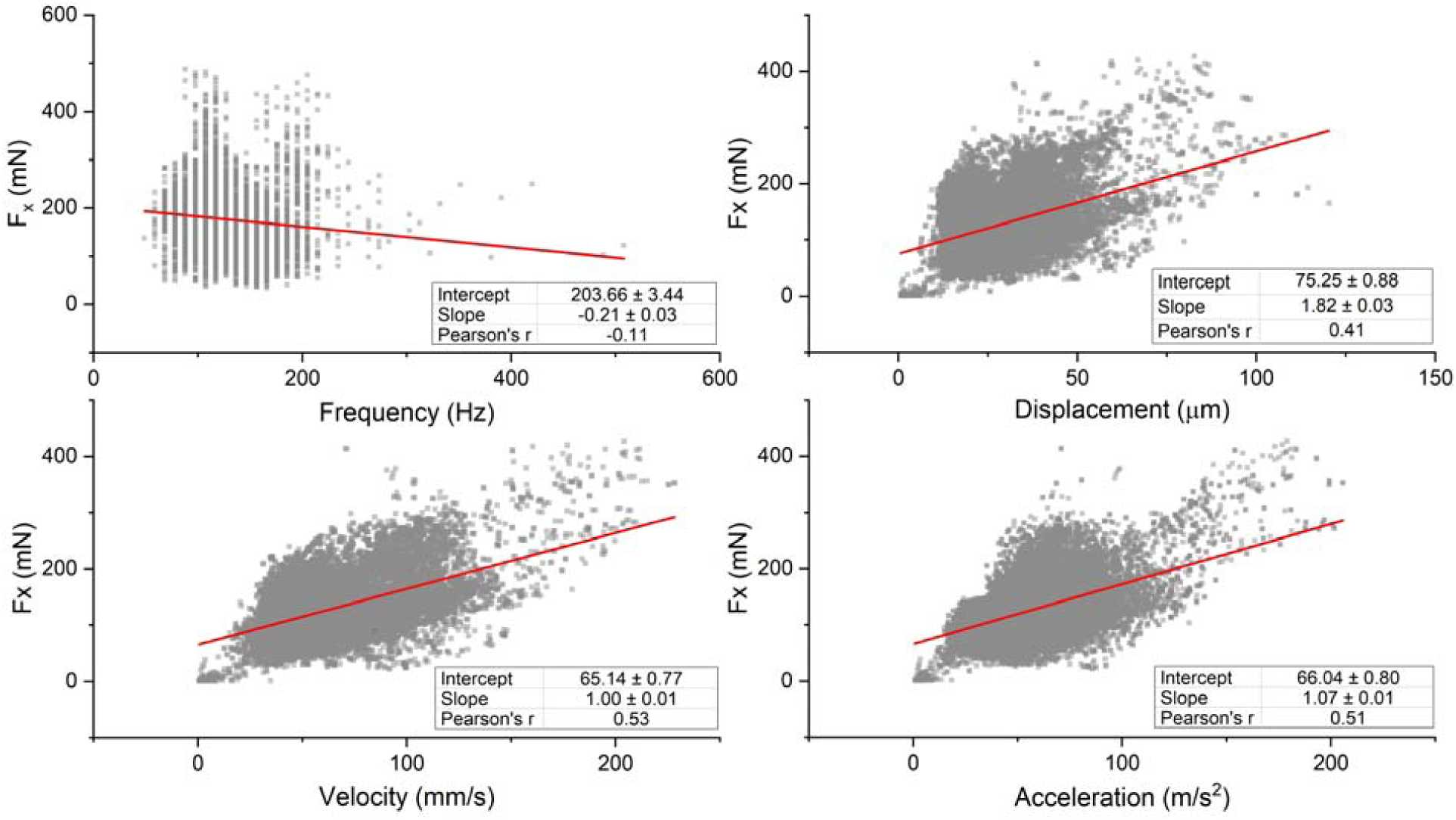
Scatter plots of thorax vibration frequency and peak displacement, velocity, and acceleration against peak force in the x-direction for all measurement intervals. The linear regression is shown in red. All regressions are significant at p < 0.001.

Linear models for individual bees are shown in Table 2. The models relating thorax kinematics to force amplitudes for individual bees have higher Pearson correlation coefficients compared to the population models in almost all cases, which may suggest some anatomical variation between the animals tested. Velocity and acceleration amplitudes have slightly higher correlation coefficients compared to displacement, indicating that velocity and acceleration amplitudes are better predictors of peak force.

**Table 2.** Linear models relating peak thorax displacement/velocity/acceleration to peak force in the x-direction. ‘Total’ indicates linear model for all measurement intervals for all carpenter bee individuals. Square brackets in the ‘total’ row indicate the 95% confidence intervals of the linear regression.

| | Displacement ($\mu\text{m}$) | | | Velocity (mm/s) | | | Acceleration ($\text{m/s}^2$) | | |
| --- | --- | --- | --- | --- | --- | --- | --- | --- | --- |
| Bee | Slope | Intercept | r | Slope | Intercept | r | Slope | Intercept | r |
| 1 | 2.01 | 76.4 | 0.59 | 1.19 | 42.6 | 0.77 | 1.35 | 44.0 | 0.70 |
| 2 | 1.03 | 107.2 | 0.59 | 1.031 | 107.2 | 0.59 | 1.27 | 82.0 | 0.72 |
| 3 | 1.42 | 92.0 | 0.81 | 1.42 | 92.0 | 0.81 | 1.21 | 133.7 | 0.72 |
| 4 | 1.88 | 41.6 | 0.41 | 1.88 | 41.6 | 0.41 | 1.83 | 34.8 | 0.56 |
| 5 | 0.88 | 9.8 | 0.62 | 0.87 | 9.8 | 0.62 | 1.38 | -0.2 | 0.72 |
| 6 | 1.02 | 15.5 | 0.79 | 1.02 | 15.5 | 0.79 | 1.27 | -9.6 | 0.81 |
| 7 | 3.91 | -8.9 | 0.52 | 1.49 | 8.5 | 0.68 | 1.33 | 18.6 | 0.57 |
| **8** | 1.61 | 46.8 | 0.62 | 1.31 | -12.1 | 0.85 | 1.29 | 6.8 | 0.85 |
| **9** | 1.85 | 54.3 | 0.42 | 1.44 | -6.3 | 0.67 | 1.45 | 13.8 | 0.57 |
| **10** | -0.24 | 117.8 | -0.08 | 1.29 | 52.1 | 0.54 | 0.635 | 88.7 | 0.46 |
| **Total** | 1.82<br>[1.77, 1.87] | 75.3<br>[73.5, 76.9] | 0.41 | 1.00<br>[0.97, 1.02] | 65.1<br>[63.6, 66.6] | 0.53 | 1.07<br>[1.04, 1.09] | 66.0<br>[64.5, 67.6] | 0.51 |

## 4. Discussion

Floral buzzing and other non-flight behaviors regulated by the IFMs are critical to the foraging ecology and defense of many Hymenoptera. Nonetheless, the directional forces generated during these behaviors, and how they relate to the kinematics of the thorax, are not well understood. In this study, we developed an experimental approach to simultaneously measure thorax vibration and the directional forces produced during defensive buzzing in *X. californica*, a behavior that we assume serves as a reasonable proxy for floral buzzing. Here, we discuss the implications of our findings within the context of floral buzzing.

We hypothesize that forces applied by the bees are large enough in amplitude to dislodge pollen from the interior locule walls and almost immediately eject pollen from the anther pores. Peak forces generated by carpenter bees averaged about 170 mN but approached 500 mN in some cases. These peak forces are approximately 30 – 80 times greater than the body weight of the insects considered (6.1 ± 0.81 mN, n = 10). Because force magnitudes are positively correlated with displacement/velocity/acceleration amplitudes (Figure 6), and velocity/acceleration amplitudes are greater during defensive buzzing than flight for other bee species [19], the forces produced during floral buzzing may be even larger than those reported here for defensive buzzing. Application of large forces to small anthers, which often weigh fractions of millinewtons [29] and have relatively low flexural rigidity [30], would cause them to deform considerably.

In addition to large force magnitudes, carpenter bees may buzz at frequencies that maximize anther deformation as well. Exciting a structure near one of its natural frequencies evokes a large vibration amplification factor [12], and thus reduces the applied force threshold required to excite the anther to the point of pollen release. Buzzing frequencies averaged about 130 Hz and were consistent across the population (Figure 4). Stamen natural frequencies (where the stamen includes a narrow filament supporting the poricidal anther) have been reported from about 45 – 300 Hz across several species of *Solanum* flowers [29]. That measured buzzing frequencies fall within the range of stamen natural frequencies suggests that carpenter bees may leverage vibration amplification to increase anther deformation and possible the amount of pollen ejected.

Furthermore, our results suggest that carpenter bees may be able to adjust their buzzing frequencies to maintain a high degree of vibration amplification between anthers. Though average buzzing frequencies were similar across the population, all individuals had the ability to modulate their buzzing frequency considerably (Figure 4). This implies that, within a range, the insect could modulate its buzzing frequency to coincide with the natural frequency of the specific anther. Indeed, bumblebees have been shown to adjust their excitation frequency based on the biophysical characteristics of the flower type they are pollinating [31]. In addition to modulating their primary buzzing frequency, carpenter bees extend their excitation bandwidth by delivering forces as repetitive impulses rather than harmonically. Repeated impulses have frequency content that begin at the primary buzzing frequency and extend to integer harmonics thereof (Figure 3). During floral buzzing, force harmonics may excite anthers with resonant frequencies higher or lower than the insect’s predominant buzzing frequency where most vibration energy occurs.

We also hypothesize that carpenter bees apply forces in a purposeful direction intended to maximize anther deformation. Previous studies indicate that *Solanum* anthers are more sensitive to vibrational forces acting in specific directions [13], [14]. The directional sensitivity of the anther can be described by the vibration modes of the anther, where a vibration mode is the spatial vibratory pattern that a structure follows if excited at one of its natural frequencies. Recent numerical studies suggest that bees excite a specific axial-bending type vibration mode during floral buzzing (Figure 7) [26]. To excite this mode, bees should apply forces in the direction and at the location where the modal displacement is highest. Bees produced the largest forces in the x and z directions (Figure 2), and usually applied forces to the anther at a location about ¾ from its tip (though to our knowledge, this bite location has not been rigorously quantified). Figure 7 resolves the axial-bending mode into x and z modal displacements (for detail on the anther structural used to estimate this vibration mode, please refer to [26]). Interestingly, this axial-bending mode has large modal displacements in both x and z directions at the location bees typically bite. Thus, the predominant forcing directions observed in this study further support the hypothesis that bees leverage vibration amplification during floral buzzing on *Solanum* anthers.

**Figure 7.**
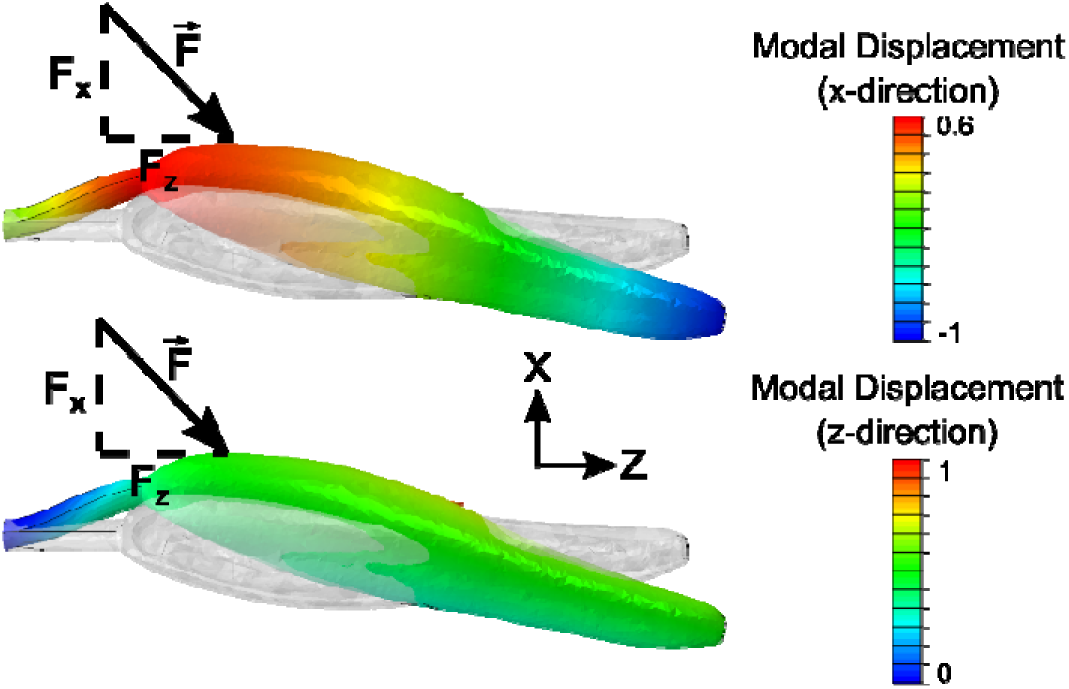
We hypothesize that bees excite this axial-bending mode during floral buzzing. The top and bottom figures show the modal displacement of the axial-bending mode in the x and z directions respectively. The force vector indicates the approximate direction and location a bee may apply forces to the anther. Note that modal displacements are arbitrary in units, and only indicate how points on a structure move in relation to one another if that mode is excited.

Lastly, we turn our attention to the relationship between thorax kinematics and force generation. Force amplitude was only weakly correlated with buzzing frequency, but was moderately positively correlated with displacement, velocity, and acceleration amplitudes (Figure 6). Thus, to increase the deformation of their thorax, the pollen foraging bee should increase the forces acting on the thorax walls. In general, the buzzing frequencies and thorax vibration velocity and acceleration amplitudes are typically 20% – 30% higher during floral buzzing than they are during defensive buzzing in bumble bees [19]. Assuming the same linear model holds for both defensive and floral buzzing behaviors, an assumption that must be tested, we would expect that the forces generated during floral buzzing would be approximately 20% – 30% higher than those measured here.

However, linear models relating thorax kinematics to force amplitudes usually had higher correlation coefficients for individual bees than the total population, in part because of differences in slopes and intercepts between individual bee models (Table 2). Variation in model slopes may arise from natural variation between individual bees. The stiffness of the thorax, for example, varies between individual honeybees [17] and individual hawkmoths [32]. If carpenter bee thoraxes have similar stiffness variability, stiffer thoraxes will require higher forces to deform during flight or other behaviors. Owing to variation between individuals, the population-wide linear models presented in this work are most appropriate for approximating the order of magnitude of peak forces for carpenter bees when displacement, velocity or acceleration magnitudes are known. Further research should be conducted to determine if our linear model is applicable to other floral buzzing species, or if other regression variables (e.g., insect mass or intertegular distance) should also be considered.

## Supporting information

Statistical Data

## Author Contributions

M.J.: conceptualization, data curation, formal analysis, investigation, methodology, project administration, writing - original draft, writing - review & editing; S.B.: conceptualization, resources, writing - review & editing; C.C.: methodology, investigation; K.B.: resources, writing - review & editing; C.H.: methodology, visualization, writing - review & editing

## Data Availability

Processed data is available at https://zenodo.org/record/5533016#.YVJbp5rMKUk.

