## Supplementary material for "Carpenter Bee Thorax Vibration and Force Generation Inform Pollen Release Mechanisms During Floral Buzzing": Statistical Data

**SUPPLEMENTARY DATA**

**Influence of Post Type**

Tests of the null hypothesis that post does not affect Fx, Fy, Fz, Displacement, Velocity, or Acceleration

**ln(Fx)**

| **Term** | **DF Numerator** | **DF Denominator** | **F-value** | **p-value** |
| --- | --- | --- | --- | --- |
| Bee (random factor) | n/a | n/a | n/a | 0.023 |
| Post | 1 | 8.01 | 0.00 | 0.978 |

**ln(Fy)**

| **Term** | **DF Numerator** | **DF Denominator** | **F-value** | **p-value** |
| --- | --- | --- | --- | --- |
| Bee (random factor) | n/a | n/a | n/a | 0.025 |
| Post | 1 | 8.00 | 1.49 | 0.257 |

**ln(Fz)**

| **Term** | **DF Numerator** | **DF Denominator** | **F-value** | **p-value** |
| --- | --- | --- | --- | --- |
| Bee (random factor) | n/a | n/a | n/a | 0.023 |
| Post | 1 | 8.00 | 0.21 | 0.660 |

**ln(Displacement)**

| **Term** | **DF Numerator** | **DF Denominator** | **F-value** | **p-value** |
| --- | --- | --- | --- | --- |
| Bee (random factor) | n/a | n/a | n/a | 0.024 |
| Post | 1 | 7.97 | 0.48 | 0.510 |

**ln(Velocity)**

| **Term** | **DF Numerator** | **DF Denominator** | **F-value** | **p-value** |
| --- | --- | --- | --- | --- |
| Bee (random factor) | n/a | n/a | n/a | 0.023 |
| Post | 1 | 8.02 | 0.84 | 0.386 |

**ln(Acceleration)**

| **Term** | **DF Numerator** | **DF Denominator** | **F-value** | **p-value** |
| --- | --- | --- | --- | --- |
| Bee (random factor) | n/a | n/a | n/a | 0.023 |
| Post | 1 | 8.06 | 1.02 | 0.341 |

**ln(Power)**

| **Term** | **DF Numerator** | **DF Denominator** | **F-value** | **p-value** |
| --- | --- | --- | --- | --- |
| Bee (random factor) | n/a | n/a | n/a | 0.023 |
| Post | 1 | 7.98 | 0.12 | 0.741 |

**Force Directionality**

Test of the null hypothesis that Fx = Fy = Fz (Figure 5)

| **Term** | **DF Numerator** | **DF Denominator** | **F-value** | **p-value** |
| --- | --- | --- | --- | --- |
| Bee (random factor) | n/a | n/a | n/a | 0.055 |
| Direction | 2 | 16 | 43.57 | <0.001 |
| Post | 1 | 8 | 0.45 | 0.520 |
| Direction * Post | 2 | 16 | 0.73 | 0.499 |
